# HEXOKINASE 1 Control of Post-Germinative Seedling Growth

**DOI:** 10.1101/548990

**Authors:** Matthew J. Lincoln, Ashwin Ganpudi, Andrés Romanowski, Karen J. Halliday

**Author notes:** **Corresponding author:** Karen J. Halliday, Institute for Molecular Plant Sciences, School of Biological Sciences, University of Edinburgh, Edinburgh EH9 3BF, United Kingdom. Matthew J. Lincoln, Ashwin Ganpudi contributed equally to this manuscript.

## Abstract

- In darkness, PHYTOCHROME INTERACTING FACTOR (PIF)-induced skotomorphogenic seedling growth, is exemplified by increased hypocotyl elongation. HEXOKINASE1 (HXK1), which is also implicated in seedling establishment, can operate as a glycolytic enzyme or as a glucose-activated sensor signalling molecule. Under light and nutrient limiting conditions, the HXK1 sensor-signalling has been shown to control hypocotyl elongation. Little is known of whether HXK1 glycolytic function, or HXK1 and PIF cross-talk, is required to control hypocotyl growth.
- We demonstrate HXK1 glycolytic activity is critical for cell expansion, and hypocotyl growth, post-germination. Notably, application of glucose-6-phosphate, the HXK1 enzymatic product, can restore short *gin2-1/hxk1-1* mutant hypocotyls to wild-type length. Further, HXK1 sensor-signalling complex components, VHA-B1 and RPT5B, do not contribute to this response, for unlike *gin2-1/hxk1-1*, the *vha-B1* and *rpt5b* alleles only disrupt hypocotyl growth following exogenous glucose application.
- mRNA-seq analysis illustrates that HXK1 and PIF signalling converge at genes with known roles in light signalling. HXK1 imposes strong regulation on chloroplast and mitochondrial encoded genes, also branched chain amino acid catabolism pathway genes, which can provide a source of respiratory substrates in starvation conditions.
- Our study establishes the importance of HXK1 enzymatic function in supporting cell expansion and hypocotyl growth. We demonstrate a degree of cross-talk between HXK1 and PIFs through common target gene set.

## Introduction

The first days of life are critical for plant survival; successful seedling establishment relies on the mobilisation of seed reserves and the perception of light through wavelength-specific photoreceptors to support emergence before the switch to photo-autotrophic growth. HEXOKINASE 1 (HXK1), a metabolic enzyme with glucose-sensor signalling properties, and PHYTOCHROME-INTERACTING FACTORS (PIFs), play a pivotal role in coupling carbon availability to growth in the developing seedling (Rolland et al, 2002, Moore et al, 2003, Stewart et al, 2011, Lilley et al, 2012, Sairanen et al, 2013, de Wit et al, 2018, Krahmer et al, 2021).

In Arabidopsis and other oilseed plants, carbon resources are stored primarily in the form of triacylglycerols (TAGs) and seed storage proteins (SSP) which account for approximately 30-40% of the seed’s dry weight each (Baud et al, 2008). After germination TAGs are degraded and converted to sucrose. TAG-derived fatty acids undergo B-oxidation, forming acetyl CoA, which fuels the production of sucrose via the glyoxylate cycle and gluconeogenesis. Sucrose is subsequently transported throughout the seedling to support development (Eastmond et al, 2000, Yang and Benning, 2018). In sink tissues, sucrose is cleaved by either SUCROSE SYNTHASE to produce UDP-glucose and fructose, or by INVERTASE to liberate glucose and fructose (Yoon et al, 2021). A vital step in the metabolism of these hexose sugars is their phosphorylation, which is mediated by HXK and FRUCTOKINASE (FRK). HXK can phosphorylate both glucose and fructose, while FRK primarily phosphorylates fructose. That said, FRK has a higher binding affinity for fructose than HXK, suggesting the principal role of HXK is the phosphorylation of glucose to glucose-6-phosphate (G6P) (Harrington and Bush, 2003, Claeyssen and Rivoal, 2007, Granot, 2007). In Arabidopsis, there are six different members of the HXK family: HXK1-3, and HEXOKINASE-LIKE (HKL) 1-3 (Karve et al, 2008). A key difference between the HXK and HKLs is the lack of glucokinase activity in the three HKL proteins, likely attributable to an indel at the adenosine binding site. Across the family, the most well studied member is HXK1.

In addition to its enzymatic role, HXK1 is known to operate as a glucose sensor-signalling molecule, directly coupling nuclear control of gene expression to glucose availability. Here, HXK1 forms a complex with partner proteins VACUOLAR H^+^ ATPASE SUBUNIT B1 (VHA-B1) and REGULATORY PARTICLE 5B (RPT5B), one of two 19S Proteasome AAA-ATPases. Chromatin immunoprecipitation (ChIP) assays indicate VHA-B1- and RPT5B-dependent HXK1, enrichment at the 5’-272 BP region of the *CHLOROPHYLL A/B BINDING 2 (CAB2)* promoter (which contains DtRE, CUF-1, CCA1, and CGF-1 binding motifs). As none of the complex proteins possess DNA-binding domains, VHA-B1 and RPT5B are proposed to interact with transcription factors to regulate gene expression (Cho et al, 2006). Genetic analysis has shown that HXK1, VHA-B1 or RPT5B mediate glucose-induced repression of the photosynthetic genes, *CAB2* and *CARBONIC*

*ANHYDRASE 1 (CAA)* expression, and glucose-induced seedling developmental arrest (Cho et al, 2006). The retention of these responses in the HXK1 catalytic mutants G104D (Gly^104^ to Asp^104^) and S177A (Ser^177^ to Ala^177^), which retain the glucose-binding site, suggested in these instances, HXK1 operated through glucose-activated signalling, rather than through its glycolytic mode (Moore et al, 2003, Feng et al, 2015).

Interestingly, HXK1 glucose-signalling is not just implicated in *CAB2/CAA* expression and developmental arrest, which are elicited by high glucose levels, but also in the regulation of hypocotyl elongation in low light and low nutrient levels (Moore et al, 2003, Cho et al, 2006). The loss of function HXK1 mutants, glucose-insensitive 2 (*gin2-1)* (Landsberg erecta) and *hxk1-3* (Columbia-0) exhibit significantly reduced hypocotyl sizes, particularly in limited light conditions (Jang and Sheen, 1997, Jang et al, 1997, Kelly et al, 2021). Additionally, the *vha-B1* and *rpt5b* mutants phenocopy the *gin2-1* short hypocotyl phenotype, while the catalytically inactive alleles *hxk1*^*G104D*^ and *hxk1*^*S177A*^ restore the WT hypocotyl response (Moore et al, 2003; Cho et al, 2006). It would be interesting to establish whether HXK1 catalytic action assists with the metabolism of seed reserves that fuel seedling growth prior to the establishment of photosynthetic competence.

The regulation of hypocotyl elongation in darkness and low light conditions is known to be regulated by PHYTOCHROME INTERACTION FACTORS (PIFs), a family of basic helix-loop-helix (bHLH) transcription factors. PIFs (PIF1, 2-8) are regulated by the phytochrome (phy) family of photoreceptors, of which phyB has been seen to be critically important in regulating seedling de-etiolation (Franklin and Quail, 2010). Excitation of the phyB chromophore with red (600-700 nm) light, induces a conformational change from the inactive (Pr) to the active (Pfr) form, which preferentially binds to PIF. This physical interaction results in rapid phosphorylation and subsequent degradation of PIFs 1, 3, 4, and 5, as well as sequestering them from their target promoters (Leivar et al, 2008, Park et al, 2012, Park et al, 2018). As a result, these PIFs are most active in darkness and low light, evident in the short hypocotyl phenotype of the quadruple *pifQ* mutant, which comprises *pif1-1, pif3-1, pif4-2*, and *pif5-3* alleles (Leivar et al, 2008). Thus, light-activated phyB repression of PIFs is critical for the switch from skotomorphogenic to photomorphogenic growth.

Both phy and PIFs have been implicated in carbon resource partitioning, where they have an important role in adjusting growth allocations to sink tissues in darkness and shaded conditions (Yang et al, 2016, de Wit et al, 2018, Krahmer et al, 2021). PIFs are involved sucrose regulation of hypocotyl growth; the *pifQ* mutant is insensitive to sucrose supplementation, and PIF5 protein stability is moderated by sucrose. (Liu et al, 2011). PIF4 has an important role in coupling carbon status to temperature-dependent hypocotyl growth (Hwang et al, 2019, Bian et al, 2022). Here, Trehalose-6-phosphate (Tre6P), a key indicator of sucrose status, stabilises PIF4 by inhibiting SNF1-RELATED PROTEIN KINASE1 (SnRK1)-mediated PIF4 phosphorylation and 26S proteasome destruction. Further, phys, together with the nutrient-sensing pathways have been shown to have a critical role in stimulating seedling shoot apical meristem stem cell division, by activating TARGET OF RAPAMYCIN (TOR) (Pfeiffer et al, 2016). More recently, a link has been made between guard cell-located PIF4 and HXK1 and sucrose-mediated hypocotyl elongation in long days. This work demonstrated that HXK1 overexpression in guard cells enhances *PIF4* expression and hypocotyl elongation in response to blue light wavelengths (Kelly et al, 2021).

In this study, we assessed seedlings under darkness and low light regimes to determine HXK1 contribution to hypocotyl elongation fuelled by seed reserve utilisation. Genetic, pharmacological, and molecular analysis point to an important role for HXK1 enzymatic function in supporting hypocotyl growth in light-limiting conditions. Our mRNA-seq data reveals HXK1 exerts wide-ranging control of metabolic genes, including a branched chain amino-acid pathway, which provides an alternative source of carbon in starvation conditions. The mRNA-seq analysis also identifies cross-talk between HXK1 and PIFs at key light signalling components. Finally, we demonstrate that several aspects of the HXK1 sensor-signalling pathway do not operate in light-limiting conditions.

## Materials and Methods

### Plant material, growth conditions, and treatments

The wild-type Arabidopsis thaliana accessions used in this study are Landsberg erecta (Ler) and Columbia-0 (Col). Seeds for mutants *gin2-1* (Ler), *hxk1-3* (Col), *pfk1-1* (Col), *vha-B1* (Col, SALK_028728), *rpt5b* (Col, SALK_069366), and *king1* (Col, At3g48530) were obtained from The Nottingham Arabidopsis Stock Centre (NASC), UK. The *hxk1*^*S177A*^ seeds were kindly shared by Professor Ruth Stadler of the University of Erlangen-Nuernberg. For all experiments, seeds were surface sterilized with bleach and Triton X-100 sown on 0.5X MS plates (0.8% agar, pH 5.7) and stratified in darkness for 2-3 days at 4°C. All plants were grown at 18°C for 4 days after stratification unless otherwise specified.

### Seedling hypocotyl length and cotyledon area measurements

Images of seedlings laid flat on growth media were used to quantify hypocotyl length and cotyledon area using ImageJ (NIH, Maryland, USA) and Adobe Photoshop CS6 (Adobe, California, USA), respectively. For glucose-6-phosphate (G6P) (Sigma G7879), sodium pyruvate (Sigma P2256), and D-mannoheptose (Sigma M6909-1G) treatments, the required concentrations from filter sterilized stocks were added to sterilized media and seeds directly sown. Unless otherwise stated, all experiments were performed in triplicate with 40 seedlings per replicate.

### Seedling hypocotyl cell counts and length measurements

For determination of etiolated hypocotyl epidermal cell lengths/numbers, seedlings were mounted on slides with water and visualized using an Eclipse E600 Nikon DIC microscope at 20x magnification. Individual cell lengths from each section of the hypocotyl (basal, middle, upper) were measured using ImageJ and cell number per file was obtained by manually counting cells from the root emergence point to the apical meristem. Unless otherwise stated, all experiments were performed in triplicate with at least 20 seedlings per replicate.

### Glucose quantification

Seedlings were harvested in liquid nitrogen, finely ground into a powder, and ethanol extracted thrice. Glucose was then quantified from these ethanol extracts using enzymatic degradation at 340nm wavelength and normalized to material fresh weight. Unless otherwise stated, all experiments were performed in triplicate with 50 seedlings per replicate

### Gene expression analysis

For qRT-PCR experiments, seedlings harvested in liquid nitrogen were ground into fine powder. Total RNA was extracted using the RNeasy Plant Mini Kit (Qiagen) with on-column DNase digestion. cDNA synthesis was performed using the qScript cDNA SuperMix (Quanta Biosciences) as described by the manufacturer. The qRT-PCR was set up as a 10μL reaction using SYBR® Green 1 480 Lightcycler® Master (Roche) in a 384-well plate, performed with a Lightcycler 480 system (Roche). Results were analysed using the Light Cycler 480 software. The primers used in this study are listed in Supplementary Table S1. Unless otherwise stated, all experiments were performed in triplicate with 50 seedlings per replicate.

### cDNA library preparation and high throughput sequencing

Total RNA was extracted from 4-day-old etiolated Ler and *gin2-1* seedlings (biological duplicates of 50 seedlings per replicate) as described above. Samples were then sent to Edinburgh Genomics (University of Edinburgh, UK) for QC check and sequencing. Briefly, quality check of the samples was performed using Qubit with the broad range RNA kit (Thermo Fisher Scientific) and Tapestation 4200 with the RNA Screentape for eukaryotic RNA analysis (Agilent). Libraries were prepared using the TruSeq Stranded mRNA kit (Illumina) and then validated. Samples were pooled to create 4 multiplexed DNA libraries, which were paired-end sequenced on an Illumina HiSeq 4000 platform (Machine name K00166, Run number 346, flowcell AHT2HKBBXX, lanes 5 and 6). On average 26.6 million 150nt PE reads were obtained for each sample.

### Processing of mRNA sequencing reads

Raw sequence reads were trimmed with cutadapt 2.8 (Martin, 2011) with default parameters and --a set to ‘AGATCGGAAGAGC’, to eliminate adapter contamination from the PE reads. Trimmed reads were aligned against the *Arabidopsis thaliana* genome (TAIR10) with HiSat2 v2.2.1 (Kim et al, 2019) with default parameters, except in the case of the maximum intron length parameter, which was set at 5000. Count tables for the different feature levels were obtained from bam files using the ‘ASpli::readCounts()’ function of ASpli package version 2.4.0 (Mancini et al, 2021) with custom R scripts and considering the AtRTD2 transcriptome (Zhang et al, 2017). Count tables at the gene level presented a good correlation overall between replicates and samples (Table S2). Raw sequences (fastq files) used in this paper have been deposited in the ArrayExpress (Kolesnikov et al, 2015) database at EMBL-EBI (www.ebi.ac.uk/arrayexpress) under accession number E-MTAB-7654.

### Differential gene expression (DGE) analysis

DGE analysis was conducted using custom R scripts for 21,256 genes whose expression was above a minimum threshold level (10 counts in 0.7 of samples in the smallest group that express the gene). DGE was estimated using the edgeR package version 3.36.0 (Robinson et al, 2010; Lun et al, 2016), and resulting P values were adjusted using a false discovery rate (FDR) criterion. Genes with p < 0.05, FDR < 0.05 and an absolute log2 fold change > 0.58 were considered differentially expressed. Volcano plots, calculation and plot of chromosomal distributions, and UpSet plots of differentially expressed genes (DEGs) were generated using R.

### GO and KEGG metabolic pathway analysis

Gene set enrichment and KEGG pathway enrichment analysis were performed using a combination of custom-written R scripts and the clusterProfiler package (Yu et al, 2012) version 4.2.2 of Bioconductor. The 21,256 expressed genes were used as the universe gene set. All KEGG pathways of *A. thaliana* were derived from the KEGG Pathway Database (http://www.kegg.jp; Kanehisa et al, 2012). Only the terms with p□<□0.05 and FDR <0.1 were further considered. Bubble plots were generated using R.

### Statistical analysis

The statistical difference between two populations was tested by two-tailed, unpaired Student’s t-test. To compare three or more populations, a one-way analysis of variance (ANOVA) followed by Dunnett’s test (comparison against a control) was performed. All analyses were done using GraphPad Prism 7 (GraphPad Software) unless otherwise indicated.

### Accession numbers and data availability

Raw sequences (fastq files) used in this paper have been deposited in the ArrayExpress (Kolesnikov et al, 2015) database at EMBL-EBI (www.ebi.ac.uk/arrayexpress) under accession number E-MTAB-7654. All custom R scripts are available at https://github.com/aromanowski/gin2_darkness. Alternatively, they are available upon request to the authors

## Results

### HXK1 supports post-germinative hypocotyl growth in a G6P-dependent manner

The *gin2-1* mutant was previously reported to have impaired hypocotyl growth in low nutrient/low light conditions (Moore et al, 2003), which led us to examine the fluence rate in which HXK1 operates. We established that the *gin2-1* (Ler) and *hxk1-3* (Col-0) hypocotyls were shorter in darkness and very low irradiance light but were indistinguishable from the wild type at fluence rates greater than 10 µmol m^-2^ s^-1^ (Fig. 1a, b). This altered fluence rate response is similar to the quadruple *pifQ* (*pif1-1, pif3-1, pif4-2, pif5-3)* mutant, though less severe. Interestingly, *gin2-1* and *hxk1-3* seedlings also have short hypocotyls in 8h light (L), 16h dark (D) short days and in longer photoperiods (16h L, 8h D), but only when the light fluence rate is low (Fig. S1). Thus, the *gin2-1* and *hxk1-3* long hypocotyl phenotypes are evident in darkness and conditions when light availability is restricted. We also observe similar reductions in the cotyledon area when light is limited (Fig. 1c).

**Fig. 1.**
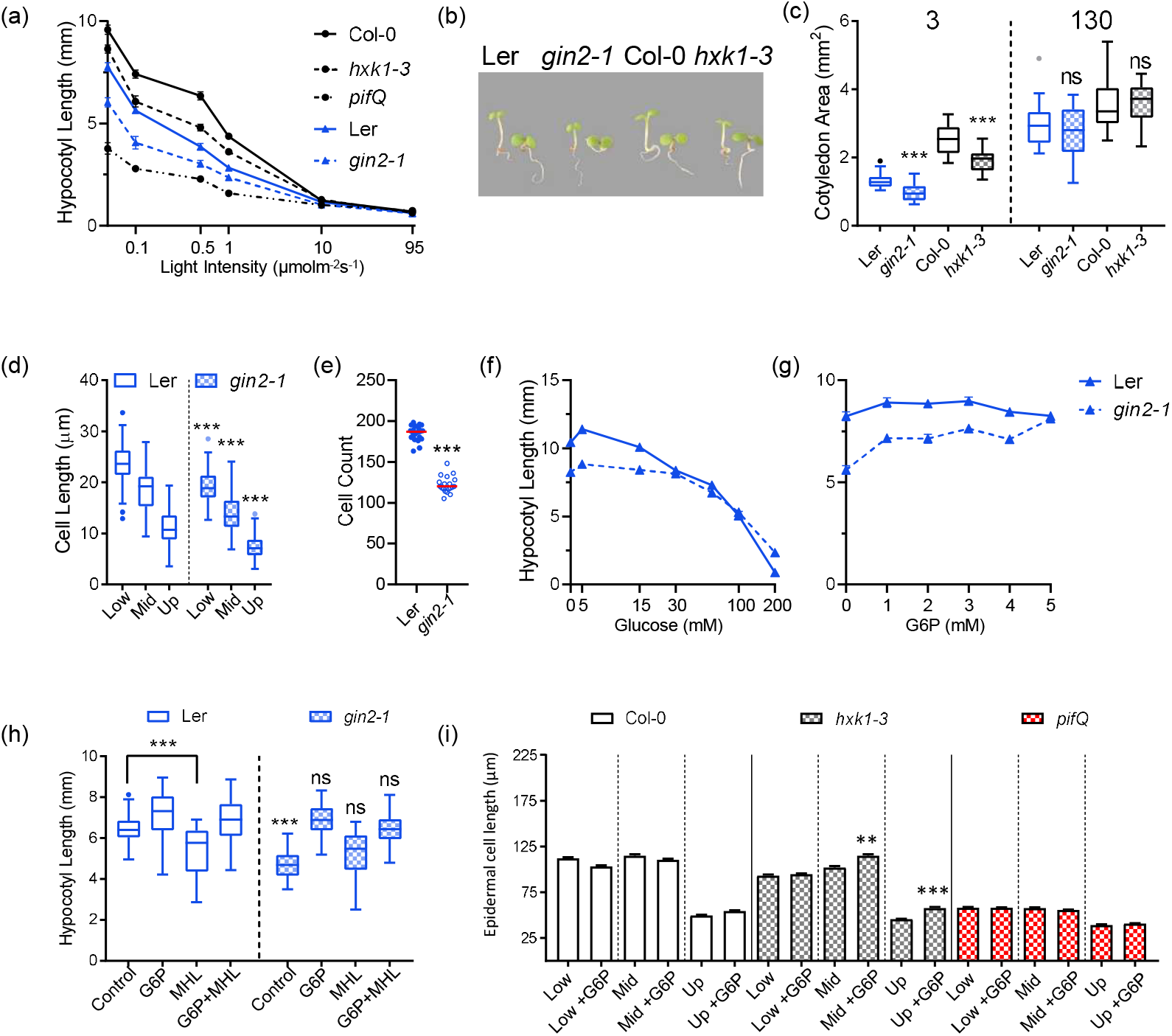
Analysis of the HXK1-dependent hypocotyl growth phenotype and dependence on G6P production (a) White light fluence response. Seedlings under continuous white light in decreasing fluences. (b) Images of seedlings. Seedlings were grown in low light (left), or high light (right). (c) Cotyledon Area. Continuous light seedlings. Numbers above represent fluence rate in m^-2^s^-1^. (d) Cell size of *gin2-1* etiolated seedlings. Etiolated seedlings. (e) Cell file of *gin2-1* etiolated seedlings. Etiolated seedlings. (f) Glucose dose-response curve. Etiolated seedlings with increasing amounts of glucose. (g) G6P dose-response curve. Etiolated seedlings with increasing amounts of G6P. (h) Competitive treatment of HXK1 inhibitor molecule mannoheptulose and glucose-6-phosphate. Short day low light seedlings without or with G6P, mannoheptulose (MHL), and both. (i) Cell size and count of *pifQ* seedlings treated with G6P. Short day low light without (left) or with 5mM G6P (right).

To establish the cellular basis of the hypocotyl defect, we measured the epidermal cell length in the basal, middle, and upper regions of the hypocotyl, and recorded the total cell number in epidermal cell files (Fig. 1d, e). Our results show that *gin2-1* has a lower cell file count and significantly shorter epidermal cells through the hypocotyl. As hypocotyl cell number is largely determined during embryo development, this suggests a role for HXK1 at this early stage. As hypocotyl elongation is driven by hypocotyl cell expansion (Gendreau et al, 1997), our data indicates that HXK1 action may be required to promote epidermal cell expansion post-germination, particularly when light conditions are limiting.

Previously, HXK1-dependent hypocotyl growth was shown to result from glucose-induced nuclear signalling, rather than HXK1 enzymatic function (Moore et al, 2003, Cho et al 2006). To explore this further, we examined the hypocotyl response over a range of glucose concentrations in low-irradiance light and in darkness. As expected, in WT the glucose response is biphasic, with lower concentrations promoting, and higher concentrations inhibiting hypocotyl growth (Singh et al, 2017) (Fig. 1f). In comparison, *gin2-1* and *hxk1-3* mutants exhibited reduced sensitivity to glucose, with a lagging doseresponse curve (Fig. 1f, Fig. S2a, Fig. S2e, S2g). This was not observed in mannitol controls in both darkness and low light (Fig. 1f, Fig. S2c-d, i-j). The results indicate that HXK1 mutants have an altered hypocotyl dose response to exogenously applied glucose. Next, we applied a concentration range of glucose-6-phosphate (G6P) which is the product of HXK1 enzymatic activity. Contrasting with the glucose response, the WT was largely insensitive to G6P. Further, we found that rising doses of G6P sequentially increased hypocotyl length in *gin2-1*, and *hxk1-3*, restoring to WT length at 5mM (Fig. 1g, Fig. S1). Similar results were obtained for seedlings supplied with competitive HXK1 inhibitory molecule, D-mannoheptose, and 5mM G6P (Fig 1h). This suggests that in light-limiting conditions, and the absence of exogenous glucose, HXK1 enzymatic activity is required to fuel hypocotyl growth. In support of this proposition, we established that G6P application to *hxk1-3* successfully restores epidermal cell length in the middle and upper regions of the hypocotyl (Fig. 1i). As post-germinative hypocotyl cell elongation proceeds in a basal to distal wave, this suggests a potential window of sensitivity to G6P (Boron and Vissenberg, 2014). Thus, in light-restricted conditions, HXK1 enzymatic activity appears to play an important role in promoting hypocotyl elongation post-germination. In comparison, hypocotyl epidermal cells of *pifQ* mutants are much shorter than WT and *hxk1-3* but are completely insensitive to G6P (Fig. 1i). This data indicates that PIF control of hypocotyl cell expansion is not due to deficiencies in G6P.

### mRNA-seq reveals a role for HXK1 in translation and carbon stress metabolism

To gain a more in-depth understanding of the HXK1 function in seedling establishment, we performed an mRNA-seq on 4-day-old etiolated Ler and *gin2-1* plants. Briefly, total RNA was extracted, and 6 stranded libraries were prepared from polyA purified RNA. These were pooled and sequenced on an Illumina HiSeq 4000 platform. Gene counts were extracted using the ASpli R package (Mancini et al, 2021) and the AtRTDv2 annotation (Zhang et al, 2017). Raw counts were filtered to remove lowly expressed genes, normalized to library size and gene expression was quantified using EdgeR (Robinson et al, 2010). This resulted in 21,256 genes (62% of the 34,212 annotated AtRTD2 genes) being considered for downstream analysis. Using custom R scripts, we performed differential gene expression (DGE) analysis using EdgeR, followed by gene ontology (GO) enrichment analysis. We discovered 2,344 genes were significantly misregulated (|logFC| > 0.58, p < 0.01 and FDR < 0.05) in *gin2-1* compared to Ler, with a similar proportion of genes down (1,156) and upregulated (1,188) (Fig. 2a, Fig. S3). When we evaluated these genes, we found that the downregulated category of genes included lipid storage, and energy-consuming processes, such as ribosome biogenesis and cell proliferation (Fig. 2b). In this figure, the bubbles represent each GO term, the size of the bubble indicating the representation factor (RF) and the colour indicating the *P-* value score. In contrast, the upregulated category had strong enrichment for genes involved in the cellular response to starvation, cellular respiration, ATP biosynthesis, amino acid catabolism and photosynthesis. These patterns are consistent with a starvation-type response and a reduced capacity to catabolise sugars.

**Fig. 2.**
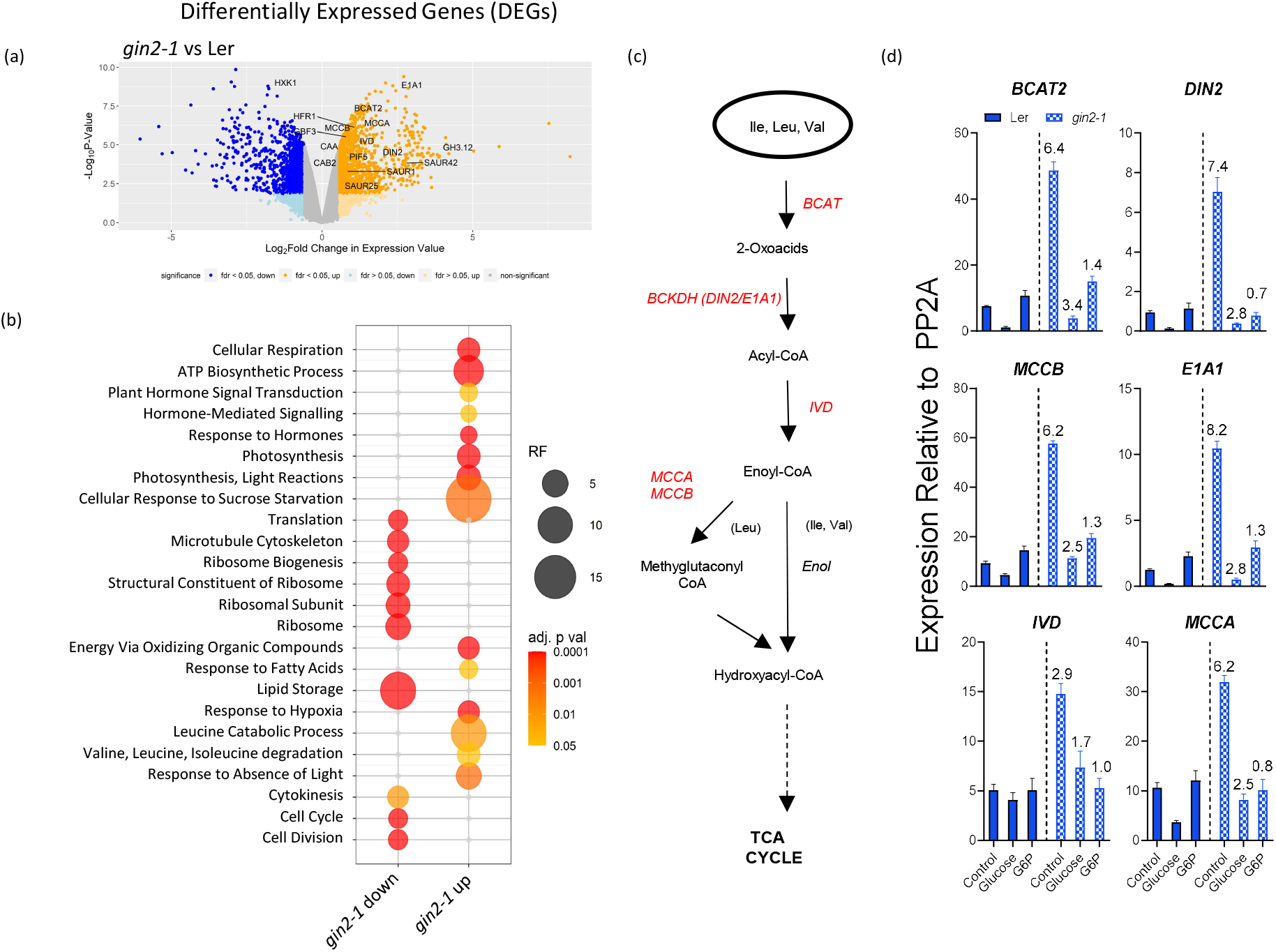
*gin2-1* sequencing data reveals importance of HXK1 in translation and carbon stress metabolism (a) mRNAseq volcano plot. Yellow points represent logFC≥0.5 and blue points represent logFC≤-0.5, pval≤0.01. Red line marks pval≤0.01. (b) Bubble plot of selected GO terms in mRNAseq. Genes of interest were collected and displayed as described in the materials and methods. (c) Diagram of branched chain amino acid metabolism in Arabidopsis. Adapted from Neinast et al, 2018 with genes represented in (d) highlighted. (d) qPCR expression of core BCAA genes misregulated in *gin2-1*. Transcript abundance of BCAA pathway genes in etiolated WT and *gin2-1* seedlings grown with or without 0.5% w/v glucose or 0.125% w/v G6P. Numbers above represent fold-change between *gin2-1* and WT.

We noted that genes that were strongly upregulated in *gin2-1* included several enzymes involved in the catabolism of the branched-chain amino acid (*BCAA*) genes: *BRANCHED-CHAIN AMINO ACID TRANSAMINASE 2* (*BCAT2), DARK INDUCIBLE 2 (DIN2)*, AT1G21400 *(E1A1), ISOVALERYL-COA-DEHYDROGENASE (IVD)*, AT1G03090 *(MCCA)*, and *3-METHYLCROTONYL-COA CARBOXYLASE* (*MCCB)* (Fig. 2c). *BCAA* genes are often suppressed in carbon-rich conditions, and induced by stress or starvation, as they provide alternative substrates for respiration (Pires et al, 2016, Binder, 2010). Indeed, we also observe strong glucose-induced suppression of the majority of these genes in both the WT and *gin2-1* (Fig. 2d). In most cases, the efficacy of glucose is greater in *gin2-1* than in WT, indicating HXK1 has a role in antagonizing the glucose response (Fig. 2c). Our data also shows that, as for hypocotyl growth (Fig. 1g), G6P application to WT has no significant impact on *BCAA* gene expression. However, G6P treatment in *gin2-1* is effective in restoring the BCAA expression to WT levels. This indicates *BCAA* gene expression is regulated by the HXK1-G6P pathway.

Another feature of the transcriptomic data analysis is the extent to which HXK1 controls organellar gene expression, with 23.5% of mitochondrial and 82.8% of chloroplast genes upregulated in *gin2-1* (Fig 3a). In the mitochondria, we observe enrichment for genes regulating metabolic proton and electron transport; and in the chloroplast, we see enrichment in all genes except for 23 genes associated with vesicle formation and chromatin remodelling. Figure 3b shows qPCR verification for the chloroplast-encoded *RNA POLYMERASE SUBUNIT* genes *RPOA, RPOB, RPOC1* and *RPOC2*, required for chloroplast gene transcription. Given the dramatic effect on the chloroplast genome, we wanted to establish whether HXK1 affected seedling greening capacity. This we tested by growing seedlings in darkness for four days and then exposing them to 3 or 100 μmol m^-2^ s^-1^ white light for 1 hour to promote chlorophyll production. We observed *gin2-1* mutants have significantly higher chlorophyll (and protochlorophyllide) levels in darkness and following both light treatments, suggesting chlorophyll production is enhanced in HXK1-deficient seedlings where resources for growth may be more limited (Fig. 3c).

**Fig. 3.**
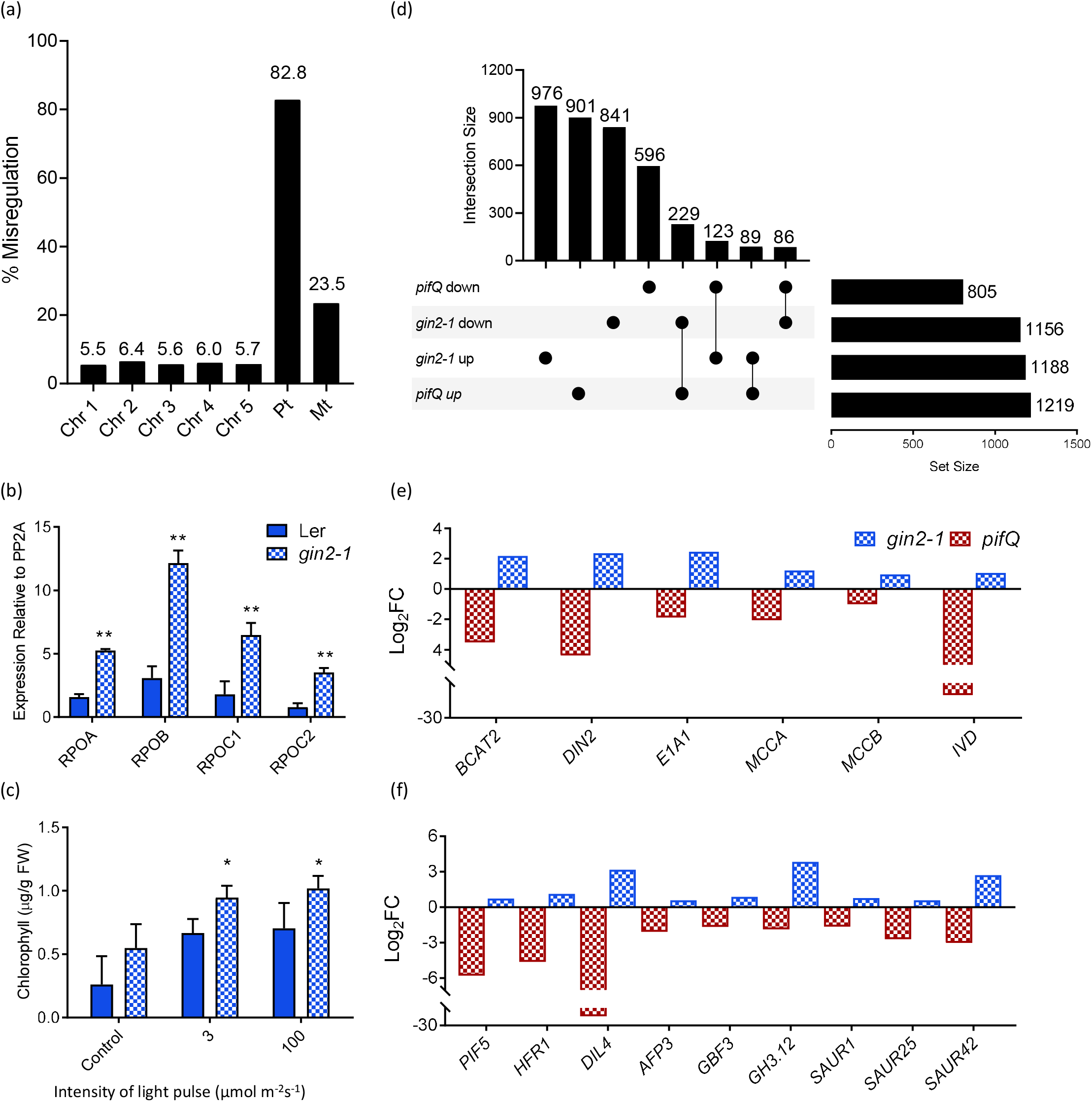
mRNAseq of *gin2-1-1* implicate a significant role in regulating the plastid genome and demonstrate crosstalk with PIF-dependent signalling (a) Percentage of chromosome misregulation in *gin2-1*. Numbers above bar represent the percentage of genes significantly (P≤ 0.01) misregulated. (b) qPCR expression of RPO family genes, collected at ZT4. (c) Chlorophyll content of *gin2-1* and WT seedlings. Seedlings grown in darkness (left) or transferred to either 3 (middle) or 100 (right)100µmol m^-2^s^-1^ white light for 1 hour. (d) Transcriptome comparison between *gin2-1* and *pifQ*. Genes significantly misregulated in *gin2-1* were compared to genes significantly misregulated in Zhang et al (2013, were categorized as being either up or downregulated in either *pifQ* or *gin2-1* and sorted into overlap and exclusive subcategories. Numbers above the bars represent count. (e) BCAA genes representation in *gin2-1* and *pifQ*. (f) Genes of interest comparison between *gin2-1* and *pifQ* with published correlations with PIF. For all figures except (c), seedlings were grown in darkness.

### Transcriptomics data Indicate HXK1 and PIF operate through distinct pathways

As HXK1 and PIFs regulate hypocotyl elongation, albeit through different mechanisms, we wanted to establish if there was evidence for pathway crosstalk or convergence. By comparing transcriptomic data from our mRNA-seq with an etiolated, 4-day-old *pifQ* mRNA-seq dataset from Zhang et al (2013), we discovered that there are 527 genes commonly misregulated between the two mutants: 175 of which are misregulated in the same direction, and 352 which are misregulated in opposite ways (Fig 3d). Interestingly, for genes that are downregulated in *gin2-1 and pifQ (gin2-1/pifQ down/down)* we observe enrichment (42 genes, or 24% of genes with synchronous regulation) in lipid storage, while in the *gin2-1*/*pifQ* up/up category, there is significant enrichment (18 genes, or 10% of genes with synchronous regulation) in hypoxia-response genes (Fig. S3). Genes with opposing *gin2-1/pifQ* down/up regulation are enriched in GO-terms groups for translation, ribosome biogenesis and cell proliferation. In contrast, opposing *gin2-1/pifQ* up/down category genes include those involved in the cellular response to starvation and amino acid catabolism. Notably, this latter group includes the *BCAA* genes *BCAT2, DIN2, THDP, IVD, MCCA* and *MCCB* that we have identified as HXK1-G6P regulated (Fig. 2c), all of which show opposing regulation by HXK1 and PIFs (Fig. 3e). We also noted that the same up/down category included a number of transcription factors such as *PHYTOCHROME INTERACTING FACTOR 5 (PIF5)* and *LONG HYPOCOTYL IN FAR-RED (HFR1)*, with pivotal roles in light signalling, *DIG-LIKE 4 (DIL4), ABA 5-BINDING PROTEIN 3 (AFP3)*, and *G-BOX BINDING FACTOR 3 (GBF3*), involved in ABA signalling, *GRETCHEN HAGEN 3*.*12 (GH3*.*12)*, the auxin conjugating enzyme, and the *SMALL AUXIN UPREGULATED RNA (SAUR)* genes *SAUR1 (AT4G34770), SAUR25 (AT4G13790)* and *SAUR42 (AT2G28085)* (Fig.3f). Thus, HXK1 has a role in repressing the expression of this group of transcription factors that are promoted by PIFs.

Our mRNA-seq data indicates that in light restricted conditions HXK1 has a role in controlling nutrient resource management, while our physiological data implicate HXK1-G6P function in stimulating hypocotyl growth. Previously mutant alleles for the *VHA-B1* and *RPT5B*, which form a nuclear complex with HXK1, were shown to restrict hypocotyl growth (Cho et al 2006). In agreement, we found *vha-B1* and *rpt5b* restricted hypocotyl elongation with glucose concentrations as low as 28mM. However, when grown on media without additional glucose supplement, *vha-B1* and *rpt5b* were indistinguishable from WT, while the *hxk1-3* phenotype was retained (Fig. 4a). We also examined the *king1 (kin*γ) mutant allele for the SNF1-RELATED PROTEIN KINASE REGULATORY SUBUNIT GAMMA 1, which was recently identified as another interaction partner for HXK1 (van Dingenen et al, 2019), though neither treatment resulted in a hypocotyl growth defect. The data suggests HXK1 can regulate hypocotyl elongation through nuclear signaling, but in the absence of exogenous glucose HXK1 enzymatic function is required.

**Fig. 4.**
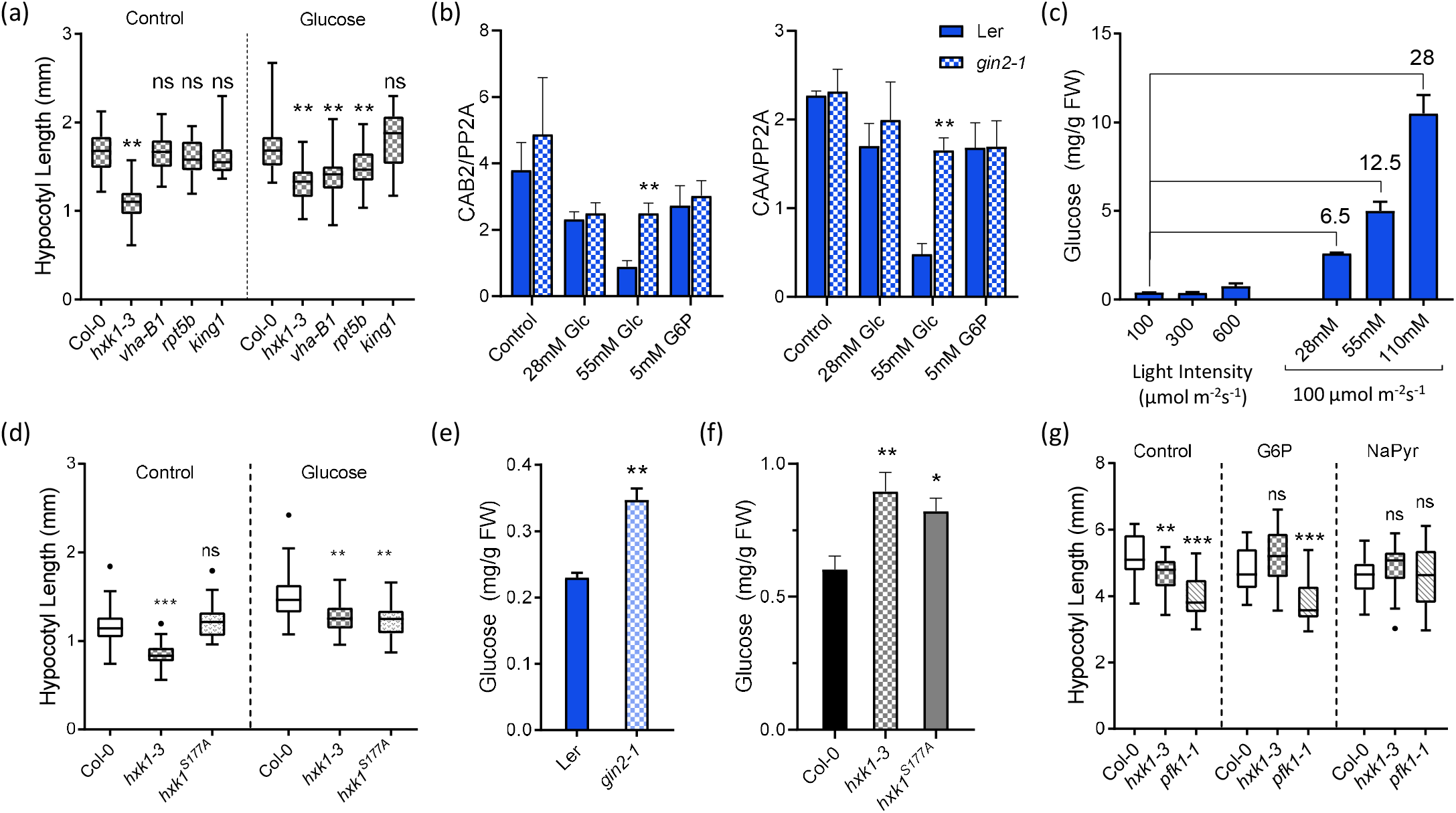
HXK1-dependent glucose signaling requires high amounts of exogenous glucose to regulate expansion and genetic regulation (a) HXK1 signaling complex mutants grown without and with additional glucose. Short day seedlings in low light, without or with 28mM glucose. (b) Expression of CAB2 and CAA in *gin2-1* with added substrate. Transcript abundance of CAB2 (left) and CAA (right) relative to PP2A concentration in Short day seedlings in high light, without or with 28mM glucose, 55mM glucose, or 5mM G6P. Plants sampled at 4 hours after dawn (ZT4). (c) Internal glucose concentrations of Wt plants under variable light and nutrient conditions. Glucose measured at ZT 4 in Wt seedlings grown in 100, 300, and 600 µmol/m^2^s, or at high light with increasing glucose. Numbers above horizontal bars represent the fold change. (d) *hxk1*^*S177A*^ seedlings grown with and without glucose. Etiolated seedlings without or with glucose. (e) Internal glucose concentration comparison between Wt and *gin2-1* seedlings. Glucose measured at ZT4 in etiolated seedlings. (f) Internal glucose concentration of *hxk1*^*S177A*^ seedlings. Glucose measured at ZT4 in etiolated seedlings. (g) Glycolysis component mutant *pfk1* grown without or with G6P or sodium pyruvate (NaPyr) in 3 µmol/m^2^s WL.

Earlier reports have implicated HXK1 (VHA-B1 and RPT5B) in glucose repression of *CAB2* and *CAA* expression, so we were interested to establish if this regulation was exclusively through the HXK1 glucose signaling route. In line with these studies, we observe HXK1-dependent repression of *CAB2/CAA* when seedlings are grown in the presence of 55mM (1% w/v), but not 28mM (0.5 % w/v) glucose (Fig. 4b). The 28mM and 55mM application doses lead to 6.5-, and 12.5-fold increases in internal when glucose levels, while a ∼115mM (2% w/v) leads to a 28-fold increase compared to controls (Fig. 4c). Further, even the lowest concentration (28mM) greatly exceeds glucose levels in seedlings exposed to the higher light fluence rates of 300 or 600 μmol m^-2^ s^-1^. We also show that *CAB2* and *CAA* expression is not substantially affected by *gin2-1* following G6P application. Our data, therefore, lends further support to the notion that *CAB2* and *CAA* are suppressed by HXK1-VHA-B1-RPT5B in response to exogenous glucose, but they are not targets of the HXK1-G6P pathway in a physiologically relevant glucose environment.

To further investigate the HXK1 enzymatic role in hypocotyl extension, we next examined the *hxk1*^*S177A*^ substitution allele, which lacks catalytic activity (Moore et al, 2003). The expectation is that if hypocotyl length (without added glucose) is due to HXK1 enzymatic activity, then *hxk1*^*S177A*^ would have a short hypocotyl phenotype. We found that on media without glucose *hxk1*^*S177A*^ hypocotyls were indistinguishable from WT, while *hxk1*^*S177A*^ hypocotyls showed reduced growth compared to WT plants when glucose is supplemented (Fig. 4d). As this finding is at odds with our other data, including the *vha-B1, rpt5B* mutant analysis (Fig 4a), we speculated that the effect of *hxk1*^*S177A*^ may be masked by a HXK1 glucose-signalling brought about by elevated internal glucose levels. Previous studies have shown that loss of HXK1 catalytic (in *gin2-1*) activity leads to over-accumulation of glucose, which we have independently verified for *gin2-1* and *hxk1-3* under our conditions (Fig. 4e). Further, this is also the case for *hxk1*^*S177A*^ that has 1.3-fold higher levels than WT (Fig. 4f). Next, we reasoned that if the *hxk1*^*S177A*^ hypocotyl elongation results from high internal glucose levels, then *hxk1*^*S177A*^ would exhibit an altered glucose dose-response. In accordance with this hypothesis, our data shows *hxk1*^*S177A*^ seedlings have an augmented dose-response that is consistent with elevated internal glucose levels (Fig. S5).

To consolidate our findings, we analysed the mutant for phosphofructokinase (PFK1), which controls a rate-limiting step in glycolysis (Mustroph et al, 2007). Our data shows that in low light the *pfk1* mutant phenocopies the *hxk1-3* short hypocotyl, and as expected *pfk1* is unresponsive to G6P but is rescued by the application of the glycolytic product, pyruvate (Fig. 4g). These results further reinforce the important role of the glycolytic pathway in supporting hypocotyl growth in light limiting conditions.

## Discussion

### HXK1-mediated hypocotyl elongation is facilitated through its glycolytic role

The seedling hypocotyl is an important model system in *Arabidopsis* for the study of signal convergence, and growth is an important readout of these signals (Oh et al, 2014, Singh et al, 2017; Ivakov et al, 2017; Chen et al, 2018). Post-germinative cell expansion requires strong coordination of these signals and is dependent upon the mobilisation of seed reserves coordinated with signalling responses (Penfield et al, 2004; Stewart et al, 2011). In this study, we have demonstrated that HXK1 plays an important role in this process via glycolytic action; through seedling establishment, HXK1 operates as a master regulator of the plastome and controls multiple aspects of the metabolic starvation response. For the most-part HXK1 operates independently of PIF signalling, though a discrete gene set including HFR1 and PIF5 were identified as possible signal convergence points.

HXK1 has been recognised to have two distinct roles: as a glycolytic enzyme which catalyses the conversion of glucose to G6P, and as a nuclear-located sugar sensor-signalling molecule (Moore et al, 2003; Cho et al, 2006). Its role in seedling hypocotyl development has previously been ascribed solely to its nuclear signalling capacity; however, our data shows that when glucose is not supplied exogenously, HXK1 glycolytic activity is necessary and sufficient to support hypocotyl growth. This becomes more apparent when light is limited, whether through a shorter light period in diurnal conditions or with reduced light fluences in both continuous light and diurnal cycles (Fig. 1a-b, Fig. S2). In contrast to glucose, where dose-response curves are qualitatively similar in WT and *gin2-1*/*hxk1-3*, G6P selectively restores the *gin2-1*/*hxk1-3* short hypocotyl to wild-type length (Fig. 1f-g, Fig. S1). These findings emphasise the importance of HXK1 in the breakdown of seed-derived carbon reserves, which fuel hypocotyl expansion in light-limiting conditions prior to activation of the photosynthetic machinery. Sucrose production from triacylglycerols and storage proteins feeds into glycolysis, which in turn allows for the production of energy, carbon skeletons and metabolites that are required for cell expansion and growth (Penfield et al, 2004, Graham, 2008). Supporting this notion, we observe HXK1-G6P mediated epidermal cell expansion in seedling hypocotyls (Fig. 1g, 1i, Fig. S1, Fig. 4g).

Earlier work has implicated PIFs in sucrose-dependent hypocotyl elongation and auxin gene expression, with PIF5 having a prominent role (Stewart et al, 2011; Lilley et al, 2012). We were therefore intrigued to establish whether PIFs had a broader role in the integration of light and HXK1 signals to control growth. Our analysis demonstrates that the short hypocotyl *pifQ* mutant is completely insensitive to G6P, suggesting that G6P is not a limiting factor for growth in *pifQ*. However, as *pifQ* only represents 4 out of the 8 *PIF* proteins in *Arabidopsis*, there is potential for other PIF-mediated growth deficiencies operating in a HXK1-G6P responsive way.

### HXK1 regulates the carbon stress transcriptome

When analysing mRNA-seq data we discovered that *gin2-1* has a transcriptome signature that is reminiscent of sucrose starvation. Earlier studies indicate sucrose starvation leads to strong upregulation of genes involved in sugar metabolism, photosynthesis, lipid metabolism, cellular respiration, and downregulation of genes involved lipid storage, cell proliferation and ribosomal processes (Contento et al, 2004, Nicolai□ et al, 2006). Our study reports a qualitatively similar pattern in the transcriptome for *gin2-1* (Fig. 2b). Amongst the *gin2-1* upregulated genes are those implicated in branched-chain amino acid (BCAA) catabolism (Fig. 2b-d), an evolutionary conserved mitochondrial pathway which can be induced by intense carbon starvation (Binder 2010, Hildebrandt et al 2015; Heinemann and Hildebrandt, 2021). Activation of the BCAA pathway provides an alternative source of respiratory substrates that can be utilised during carbohydrate starvation. As G6P supplementation specifically rescues the expression of these genes, our data indicates *BCAA* genes are regulated through HXK1-G6P induced signalling. Interestingly, the expression of *BCAA* genes is known to be directly regulated by SnRK1 kinase-bZIP complexes (Pedrotti et al, 2018). This evolutionarily conserved protein kinase complex acts to maintain homeostasis under nutrient-stress conditions. It will therefore be interesting to establish if SnRK1 kinase-bZIP complexes can be regulated by HXK1-G6P.

### In light limiting conditions HXK1 exerts strong control over the plastome

Our mRNA-seq data reveals a significant effect of HXK1 on the expression of non-nuclear genes. In dark grown seedlings 82.8% of the chloroplast and 23.5% of the mitochondrial genes were elevated in *gin2-1* (Fig. 3a-c, Table S2). While glucose treatment has been shown to attenuate chlorophyll degradation in darkness, our data show HXK1 represses chloroplast gene expression in darkness, which, in turn, suppresses chlorophyll levels following exposure to light (Ueda et al, 2020). Indeed, the connection between chloroplast biogenesis and metabolism is well established; albino mutants show high misregulation in genes involved with starvation and mobilisation of storage energy, consistent with the notion of chloroplast genesis representing a switch of growth strategies to autotrophism (Grübler et al, 2017). With respect to mitochondria, sucrose starvation has been shown to cause structural changes, which result in impaired protein production despite unimpeded transcription activity (Giegé et al, 2005). Additionally, we have shown HXK1 represses the expression of mitochondrial genes involved in processes including charged particle transport, ATP synthesis, and oxidative responses. Oxidation in the mitochondria has been linked to metabolism, feeding back into metabolic processes under both abiotic and biotic stress responses to maintain signalling capacity (Vanlerberghe, 2013). The impact of *gin2-1* on these organellar transcriptomes most likely reflects the reality of carbon-resource deprivation, and the need to prime the cellular machinery to switch to photosynthesis rather than reliance on limited seed reserves to fuel growth.

### HXK1 and PIF signalling converges at a small gene subset

Our mRNA-seq analysis identified small group of genes (22% of total *gin2-1* misregulation) that exhibit similar or opposing regulation by PIF and HXK1 (Fig. 3d-f, Fig. S4). We were interested to establish that *BCAA* genes, identified as HXK1-repressed genes (Fig 2a-d), were promoted by PIFs. Likewise, a number of genes implicated in light signalling including *PIF5, HFR1, GH3*.*12* and *SAUR* genes, were antagonistically regulated by HXK1 and PIFs. This suggests that while HXK1 and PIF-mediated signalling is largely distinct, crosstalk does occur through a few key components. Indeed, a recent study demonstrated such a connection between HXK1 and PIF4 (Kelly et al 2021). The study showed HXK1 located in guard cells controls hypocotyl elongation through PIF4, in white and blue light. The authors suggest that in contrast to red light, blue light is not effective in driving photosynthesis, and therefore in blue light conditions sucrose derived from seed storage is a major carbon source. The study has strong parallels to our low light study where hypocotyl growth is largely fuelled by seed reserves. In both instances, HXK1 has an important metabolic role in phosphorylating the sucrose cleavage products, glucose and fructose. Like HXK1, PIF transcription factors are primed to operate under low light conditions, and thus offer a route to couple carbon resource management to growth prior to the establishment of photoautotrophic growth.

### HXK1-dependent glucose signalling requires high amounts of glucose to operate

Previous work has shown *VHA-B1* and *RPT5B* form a nuclear complex with HXK1, which directly couples glucose sensing to nuclear signalling and hypocotyl growth (Cho et al, 2006). In agreement with this notion, we observe a hypocotyl defect in *vha-B1, rpt5b* mutants, but only when exogenous glucose is supplied. In contrast the *gin2-1*/*hxk1-3* short hypocotyl phenotype is observed both with and without glucose (Fig. 4a). Collectively the data indicate hypocotyl elongation can be controlled through HXK1 glycolytic and sensor-signalling pathways, however, prior to the establishment of photoautotrophic growth the glycolytic pathway necessarily dominates. These observations are backed by qPCR analysis of *CAB2* and *CAA1*, both HXK1/VHA-B1-/RPT5B-dependent glucose repressed genes. Interestingly, we only observed HXK1-mediated repression at 55mM glucose (which elevates internal glucose levels by 12.5-fold) and not the lower concentration of 28mM glucose (6.5-fold increase) (Fig. 4b). This suggests HXK1-dependency is only observed when glucose levels exceed normal physiological levels, which may be more indicative of a stress response. The ABA signalling genes *ABI4* and *ABI5* have been shown to have key roles in abiotic stress and the glucose signalling response (Dijkwel et al, 1997, Arenas-Huertero et al, 2000). As ABI4 is known to directly regulate the expression of *CAB2* it will be interesting to establish whether the HXK1-VHA-BI-RPT5B complex operates with ABI4 or other ABA signalling components.

Our study provides strong evidence for HXK1 glycolytic control of hypocotyl growth, yet counterintuitively, the *hxk1*^*S177A*^ catalytically inactive mutant does not have the expected short hypocotyl phenotype in low light (Moore et al, 2003; Fig. 4d, S4). We therefore speculated that in a similar manner to *gin2-1* and *hxk1-3*, loss of enzymatic activity could lead to enhanced internal glucose levels. This would effectively activate glucose signalling and hypocotyl elongation, which would effectively mask any growth impairment caused by loss of enzymatic function (Fig. 4e-f,). Our analysis, which revealed elevated glucose levels in *hxk1*^*S177A*^ and a shifted glucose dose-response, provided support for this hypothesis, offering a rational explanation for the *hxk1*^*S177A*^ phenotype.Corroborative evidence for the importance of glycolysis, comes from genetic analysis demonstrating the glycolytic enzyme mutant *pfk1-1* phenocopies *hxk1-3*, while both mutants are rescued by exogenous pyruvate application (Fig. 4g).

In conclusion, this study has shown HXK1 performs an important role in post-germinative hypocotyl growth, by enabling the consumption of seed reserves, prior to establishment of photosynthetic competence. By controlling the expression of enzymes in the BCAA pathway, HXK1 provides a route to generate alternative respiratory substrates when carbon resources are limited. HXK1 control of the plastome, provides a means to link photosynthetic machinery establishment to carbon needs. While HXK1 control of light signalling components provides a potential route to couple resource availability to the regulation of growth.

## Acknowledgements

M.L. and A.G. were supported by the University of Edinburgh Darwin Trust Scholarship. A.R was supported by Biotechnology and Biological Sciences Research Council (BBSRC) grant BB/N005147/1, awarded to K.J.H.)

## Author Contributions

M.J.L., A.G., A.R. and K.J.H. planned and designed the research. A.G designed and performed, and analysed the mRNA-seq experiment, A.R. conducted the bioinformatics analysis of the mRNAseq data. All other experiments were performed and analysed by M.J.L and A.G. All authors contributed to manuscript preparation.

