## Supplemental figures for "HEXOKINASE 1 Control of Post-Germinative Seedling Growth"

(a)

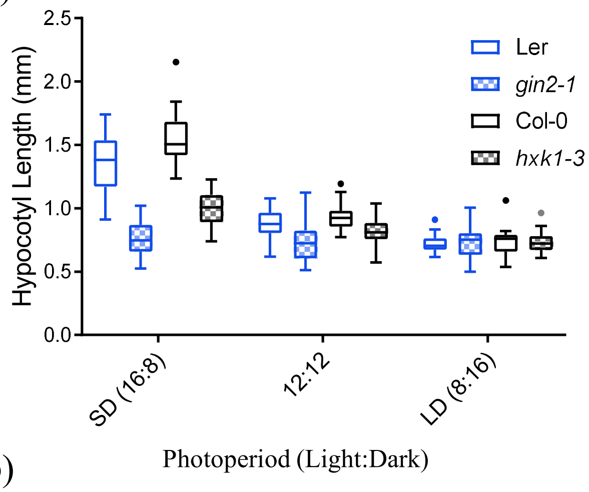


(b)

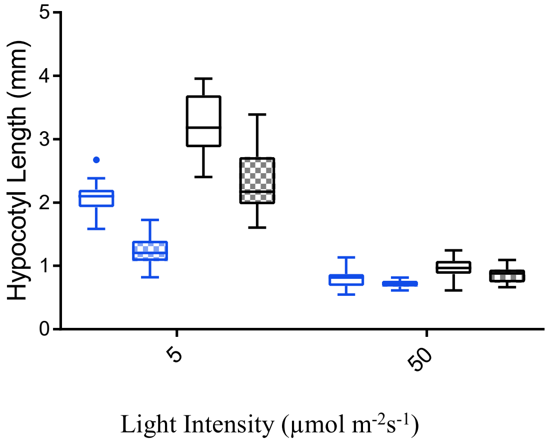


**Fig S1**. Diurnal growth of *gin2-1* and *hxk1-3*. (a) Effect of photoperiod on *hxk1* seedling growth. Diurnally grown seedlings grown in 100µmol $m^{-2}s^{-1}$ white light. (b) Effect of fluence rate on long day grown *hxk1* seedlings. Seedlings grown in long day.

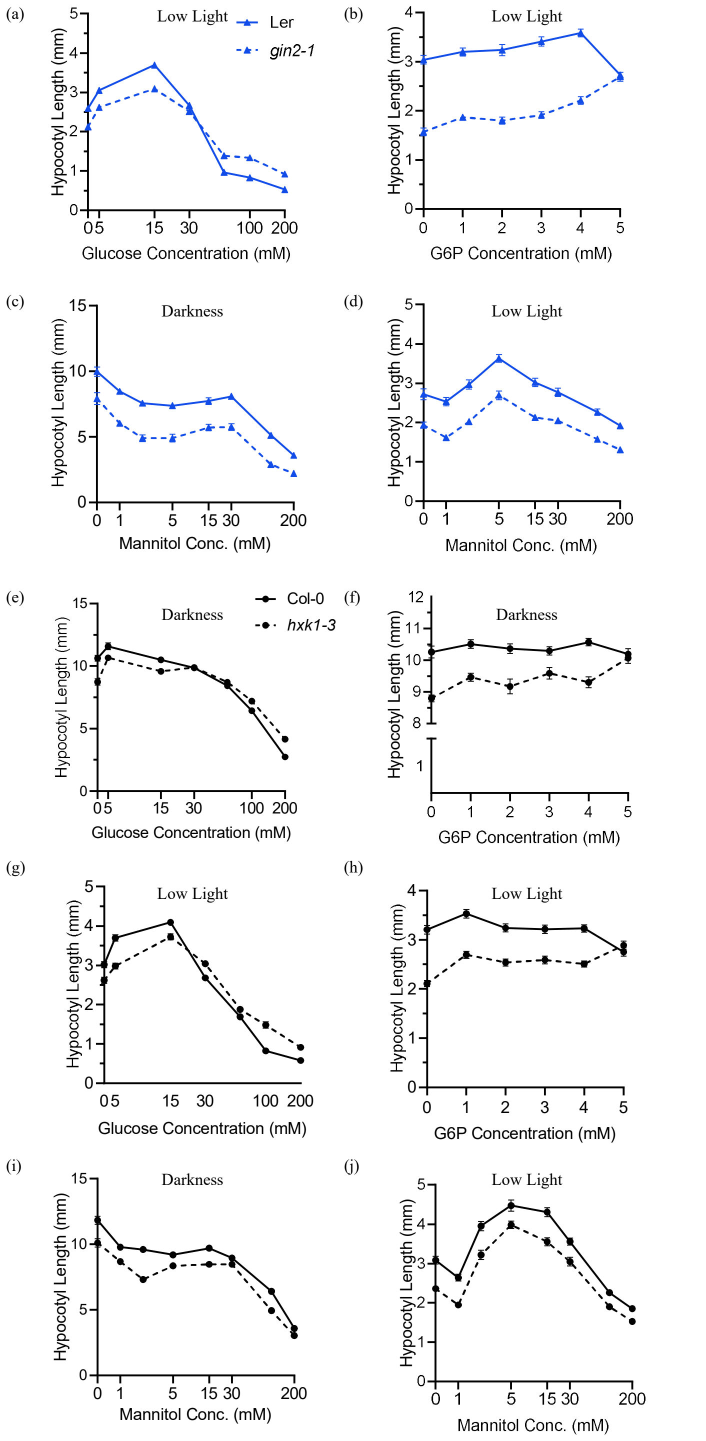


**Fig. S2** Osmotic controls and low light treatment supports importance of G6P in HXK1-dependent hypocotyl elongation. (a) Low light glucose, Ler and *gin2-1*. (b) Low light G6P, Ler and *gin2-1*. (c) Darkness mannitol, Ler and *gin2-1*. (d) Low light mannitol, Ler and *gin2-1*. (e) Darkness glucose, Col-0 and *hxk1-3*. (f) Darkness G6P, Col-0 and *hxk1-3*. (g) Low light glucose, Col-0 and *hxk1-3*. (h) Low light G6P, Col-0 and *hxk1-3*. (i) Darkness mannitol, Col-0 and *hxk1-3*. (j) Low light mannitol, Col-0 and *hxk1-3*.

| Category | Misregulated Genes | Total Expressed Genes | % Misregulation |
|---|---|---|---|
| *gin2-1* up | 1,188 | 21,256 | 5.59 |
| *gin2-1* down | 1,156 | 21,256 | 5.44 |

**Table S1.** Proportions of genes represented in the mRNAseq in **Fig. 4a**.

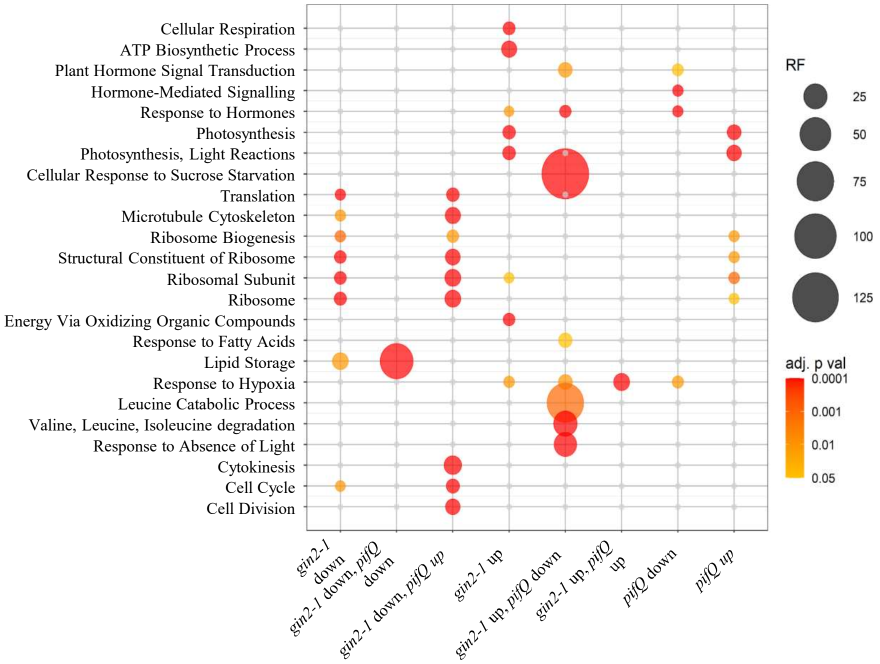


**Fig. S3** Bubble plot of selected GO terms in *gin2-1* and *pifQ* comparative analysis. Genes of interest were collected and displayed as described in the materials and methods.

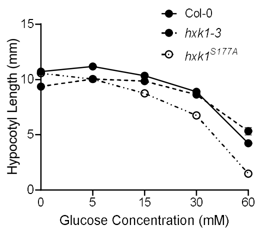


**Fig. S4** Glucose dose-response curve of *hxk1*$^{S177A}$ seedlings. Etiolated seedlings with increasing amounts of glucose.

| Primer name | Sequence (5’-3’) |
|---|---|
| AG_2250_RPOA_F | GCGATGCGAAGAGCTTTACT |
| AG_2251_RPOA_R | CCAGGACCTTGGACACAAA |
| AG_2252_RPOB_F | GATGTGAGGTGGGTTCAGAA |
| AG_2253_RPOB_R | GGTCTCCCGTCTTGCAAATA |
| AG_2254_RPOC1_F | TTCTTCCTCCCGAGTTGAGA |
| AG_2255_RPOC1_R | CCACGGCTTCTTGTACCAAT |
| AG_2256_RPOC2_F | CGCGTCGACTTGTTGAAGTA |
| AG_2257_RPOC2_R | CGTCTGCTAAGACACGACCA |
| AG_2236_BCAT2_F | GGGATAATCTCGGGTTTGGT |
| AG_2237_BCAT2_R | CTTCATCCGGATAGCGTTGT |
| AG_2232_THDP_F | GACGAAGACGGACGAATCAT |
| AG_2233_THDP_R | TGCTGAAGCGATGTTAATGG |
| AG_2234_DIN2_F | CGGTCGTCGGAGAGAGTAAC |
| AG_2235_DIN2_R | GCCTTGCAAAACACCAAAAT |
| AG_2238_MCCA_F | CCCGTCTACAGGTCGAACAT |
| AG_2239_MCCA_R | ACCCGAACTGATGGTGAGAC |
| AG_2240_IVD_F_ | ACTCTGTTGCGAGGGACTGT |
| AG_2241_IVD_R | CTTAGAAGGCGTCCTGTTGC |
| AG_2242_MCCB_F | CTTTGCCTTCAGGTGGGATA |
| AG_2243_MCCB_R | ACCGAGCAGCAATCTCTTGT |
| AG_1267_CAB2_F | CCCTGGAGACTACGGATG |
| AG_1267_CAB2_R | TCCAAACTTGACTCCGTTCC |
| AG_1428_CAA_F | TGAATACGCTGTCTTGCACC |
| AG_1429_CAA_R | TGTGATGGTGGTGGTAGCGA |
| DY_1166_HXK1_F | GGTTTCACTTTCTCGTTTCCTG |
| DY_1167_HXK1_R | CTTGTCCAACTGCTTCTTCG |
| AR027_PP2A_F | TAACGTGGCCAAAATGATGC |
| AR028_PP2A_R | GTTCTCCACAACCGCTTGGT |

**Table S2.** Primers used in the course of this study

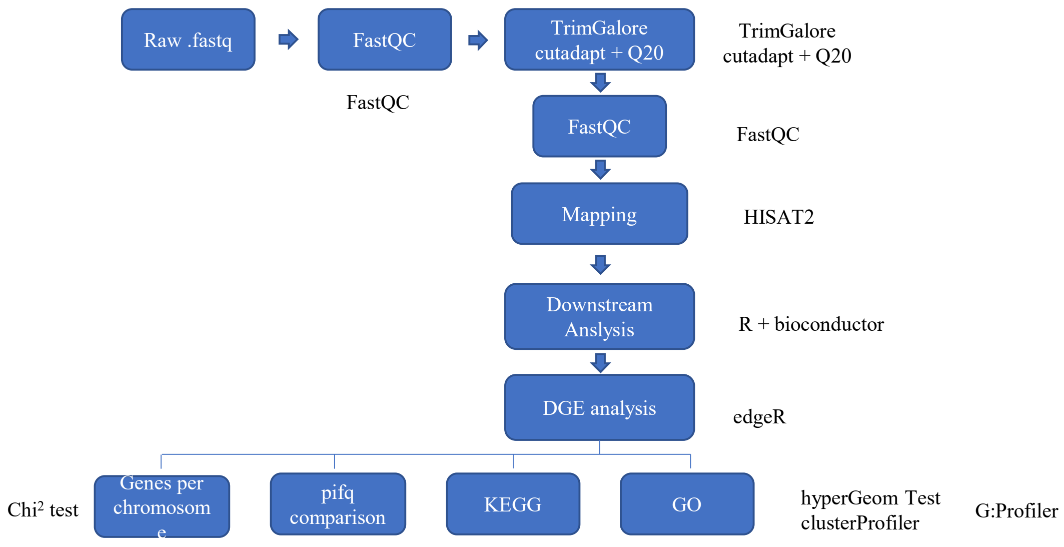


**Fig. S6** Data analysis pipeline of mRNAseq

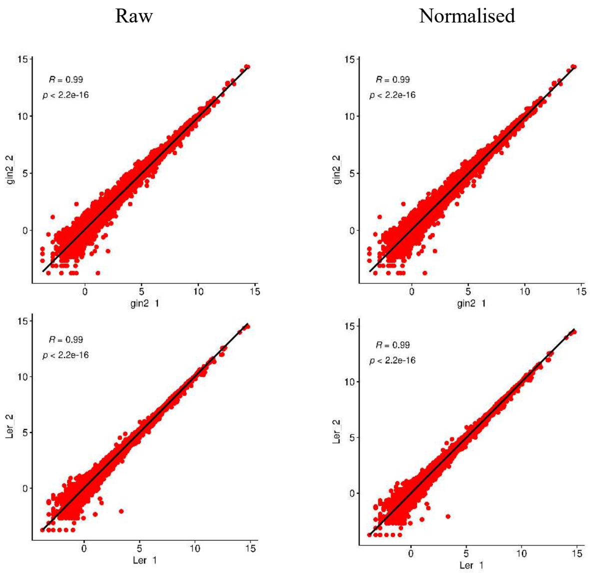


**Fig. S7** Person sample-similarity correlation of *gin2-1* and Ler samples used in mRNAseq

| Chromosome | Genes/ chromosome | Total Genes | Ratio | Total DEGs | Expected | DEGs/ chromosome | χ2 Values | Pval | $-Log_{10}p$ |
|---|---|---|---|---|---|---|---|---|---|
| 1 | 9701 | 3814 | 0.253993 | 2344 | 595.3591 | 530 | 7.18E+00 | 7.39E-03 | 2.13E+00 |
| 2 | 6312 | 3814 | 0.165262 | 2344 | 387.3731 | 407 | 9.94E-01 | 3.19E-01 | 4.97E-01 |
| 3 | 7624 | 3814 | 0.199613 | 2344 | 467.8917 | 427 | 3.57E+00 | 5.87E-02 | 1.23E+00 |
| 4 | 5842 | 3814 | 0.152956 | 2344 | 358.5288 | 353 | 8.53E-02 | 7.70E-01 | 1.13E-01 |
| 5 | 8419 | 3814 | 0.220427 | 2344 | 516.6816 | 478 | 2.90E+00 | 8.88E-02 | 1.05E+00 |
| Pt | 134 | 3814 | 0.003508 | 2344 | 8.2237 | 111 | 1.28E+03 | 2.70E-281 | 2.81E+02 |
| Mt | 162 | 3814 | 0.004242 | 2344 | 9.942085 | 38 | 7.92E+01 | 5.66E-19 | 1.82E+01 |

**Table S3.** Count tables at the gene level organised under chromosome and plastome. DEG=Differentially Expressed Genes.
